# Detecting pairwise correlations in spike trains: an objective comparison of methods and application to the study of retinal waves

**DOI:** 10.1101/006635

**Authors:** Catherine S Cutts, Stephen J Eglen

## Abstract

Correlations in neuronal spike times are thought to be key to processing in many neural systems. Many measures have been proposed to summarise these correlations and of these the correlation index is widely used and is the standard in studies of spontaneous retinal activity. We show that this measure has two undesirable properties: it is unbounded above and confounded by firing rate. We list properties needed for a measure to fairly quantify and compare correlations and we propose a novel measure of correlation — the Spike Time Tiling Coefficient. This coefficient, the correlation index and 33 other measures of correlation of spike times are blindly tested for the required properties on synthetic and experimental data. On the basis of this, we propose a measure (the Spike Time Tiling Coefficient) to replace the correlation index. To demonstrate the benefits of this measure, we re-analyse data from seven key studies which previously used the correlation index to investigate the nature of spontaneous activity. We re-analyse data from *β*2(KO) and *β*2(TG) mutants, mutants lacking connexin isoforms and also the age-dependent changes in wild type and *β*2(KO) correlations. Re-analysis of the data using the proposed measure can significantly change the conclusions. It leads to better quantification of correlations and therefore better inference from the data. We hope that the proposed measure will have wide applications, and will help clarify the role of activity in retinotopic map formation.

## Introduction

Quantifying the degree of correlation between neural spike trains is a key part of analyses of experimental data in many systems (Kirkby et al., 2013; Chiappalone et al., 2006; Dehorter et al., 2012). Neural coordination is thought to play a key role in information propagation and processing and also in self-organisation of the neural system during development. For example, correlated activity plays a critical role in forming the retinotopic map (Feller, 2009). In the developing retina, waves of correlated spontaneous activity in retinal ganglion cells have been recorded (on multielectrode arrays, MEAs, and by calcium imaging) in-vitro in many species (Wong, 1999) and shown in-vivo using calcium imaging in mouse (Ackman et al., 2012). These waves show both temporal and spatial correlations. Much work has focused on assessing the role of this activity in the development of the retinotopic map; typically both the map and various statistics of the activity are compared between wild type and mutant genotypes. The results are used to make inferences about which features of the activity are implicated in retinotopic map formation, e.g. Stafford et al. (2009). There is strong evidence that correlation between neuronal spike times is involved in this process (Xu et al., 2011).

An appropriate quantification of these correlations is vital for inference about their role. Quantifying correlations is challenging for two reasons. Firstly, correlated neurons fire at similar times but not precisely synchronously so correlation must be defined with reference to a timescale within which spikes are considered correlated. Secondly, spiking is sparse with respect to the recording’s sampling frequency (spiking rate about 1 Hz, sampling rate typically 20 kHz (Demas et al., 2003)) and also spike duration. This means that conventional approaches to correlation (such as Pearson’s correlation coefficient) are unsuitable as periods of quiescence should not count as correlated and correlations should compare spike trains over short timescales, not just instantaneously.

Many alternative measures of quantifying correlations exist (e.g. Kruskal et al. (2007); Kerschensteiner and Wong (2008); Joris et al. (2006)). One measure, the correlation index (Wong et al., 1993), has widespread popularity and is the standard measure applied to spontaneous retinal activity. It also has wider uses such as quantifying correlations in motor (Personius et al., 2007) or hippocampal (MacLaren et al., 2011) neurons. It is a pairwise measure which quantifies temporal correlations and is frequently used to investigate their dependency on a third variable, such as neuronal separation and to compare correlations across phenotypes. We show that the correlation index is confounded by firing rate which means it cannot fairly compare correlations. We list properties required of a correlation measure and conduct a thorough literature search for other measures. We propose a novel measure and then blindly and systematically test all measures for the required properties against synthetic and experimental data to propose a replacement for the correlation index. Using this replacement, we then re-analyse data from seven studies to show that results can change when correlations are measured in a way which is independent of firing rate.

## Materials and Methods

### Analysis of correlation index

The correlation index *i_A,B_* between two spike trains A and B is defined as the factor by which the firing rate of neuron A increases over its mean value if measured within a fixed window (typically 0.05–0.10 s) of spikes from B (Wong et al., 1993). The following notation is used throughout: the vectors **a** and **b** represent the spike times of neurons A and B; *a_i_* is the *i*th spike in train A and *b_j_* is the *j*th spike in train B. The correlation index is given by

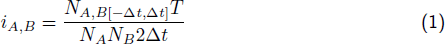

where

*N_A_*= |**a**| (total number of spikes of A in recording),

*N_B_*= |**b**|,

*T* = length of recording,

Δ*t* = synchronicity window,

and *N_A,B_*[*−*Δ*t,* Δ*t*] is the number of spike pairs where a spike from train A falls within *±*Δ*t* of a spike from train B:

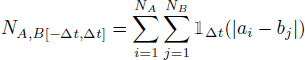

where

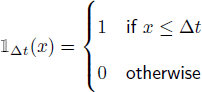

To show that the correlation index is dependent on firing rate, we assume the following model for neuronal firing: spike trains A and B are both Poisson processes with rates *λ_A_* and *λ_B_* respectively. A fixed proportion of the spike times are shared (A and B fire spikes synchronously). These synchronous spikes occur with a rate *λ_S_ ≤ λ_A_, λ_B_*. Adjusting these parameters can scale the rates whilst maintaining the correlation structure to test for rate-dependence. Under this model an expression for the correlation index can be derived:

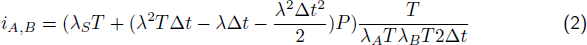

This expression is clearly rate-dependent. Two minor assumptions about the size of the rates, *T* and Δ*t* were used to arrive at this result. These assumptions are valid within experimentally observed ranges and the rate-dependency of *i_A,B_* is not affected if the assumptions are violated. In Results, we use the sub-case of auto-correlation *λ* = *λ_A_* = *λ_B_* = *λ_S_* to show that this rate-dependence significantly affects correlation values. In this case the correlation index is:

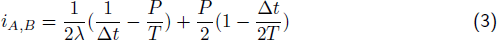

This dependency was verified computationally by extensive testing on synthetic data, including data generated from the above model using freely available code (Macke et al., 2009).

### Spike Time Tiling Coefficient

We define the Spike Time Tiling Coefficient in Figure 1. To quantify the correlation between spike trains A and B, we look for spikes in A which fall within *±*Δ*t* of a spike from B. We consider the proportion of spikes in A which have this property as this is insensitive to firing rate. We account for the amount of correlation expected by chance by making the minimal assumption that we expect the proportion of spikes from A falling within *±*Δ*t* of a spike from B *by chance* to be the same as the proportion of the total recording time which falls within *±*Δ*t* of a spike from B. Any extra spikes in A which have this property are indicative of positive correlation. We therefore use the quantity *P_A_ − T_B_* (see Figure 1 for definitions) which is positive if spikes in train A are correlated with spikes from train B, and negative if there is less correlation than expected by chance. We require the coefficient to be equal to + 1 for auto-correlation, to be *−*1 when *P_A_* = 0, *T_B_* = 1 and to have a range of [*−*1, 1]. The normalisation factor (1 *− P_A_T_B_*) ensures that these criteria hold.

**Figure 1:**
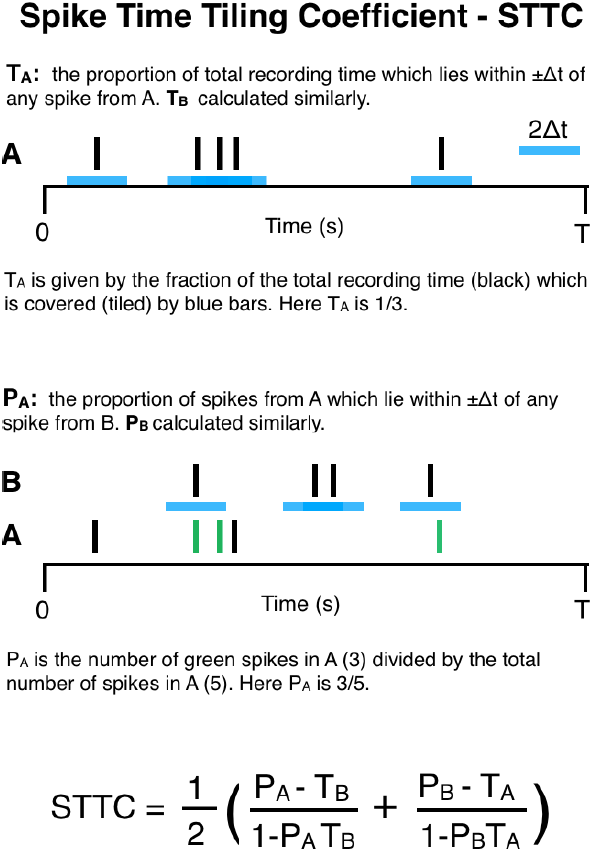
Cartoon to demonstrate the calculation of the Spike Time Tiling Coefficient. The four quantities required to calculate the Spike Time Tiling Coefficient are *P_A_*, *P_B_*, *T_A_*, *T_B_*. The only free parameter is Δ*t*. Values and scales are for demonstration only.

The coefficient should be symmetric so we consider both (*P_A_ − T_B_*) and (*P_B_ − T_A_*), combine the contributions from both trains and re-normalise to preserve the required range (see Figure 1). Computation of the Spike Time Tiling Coefficient is straightforward; the only complexity is ensuring that overlapping tiles do not count multiple times when calculating *T_A_* and *T_B_*.

The Spike Time Tiling Coefficient uses the proportion of the recording which falls within *±*Δ*t* of spikes from A to determine if the proportion of spikes in B which also have this property is indicative of correlation (i.e. more or less than is expected by chance). Since tile overlaps are not counted multiple times, this depends on the firing patterns as well as rates. In the correlation index (and many other measures) only the firing rates are used to assess what is expected by chance, but firing patterns are, in fact, important. For instance, consider an extreme case of two spike trains (A and B) with the same average firing rate where the spikes in A occur at regular intervals (no two spikes are within Δ*t* of each other) and the spikes in B all occur within Δ*t* of each other. More of the recording time lies within *±*Δ*t* of any spike from A compared to B so given an arbitrary train C, we expect more spikes in C to fall within *±*Δ*t* of any spike from A by chance than within *±*Δ*t* of any spike from B. This information is not captured using the firing rates to assess what we expect to occur by chance but is captured with the Spike Time Tiling Coefficient.

### Implementing the measures

A literature search produced 33 other measures which were implemented for testing. All measures were implemented in R and C except Victor and Purpura (1997), ISI-distance (Kreuz et al., 2007a), Van Rossum (2001) and SPIKE (Kreuz et al., 2013) which were run using freely available MatLab code (Kreuz et al., 2013). Some measures were altered to make them more likely to posses the required properties. The following changes were made (see Table 3): the Kerschensteiner and Wong (2008) and Jimbo et al. (1999) indices are originally defined as functions over binned time lags, the value of the bin around zero was taken to be the value of the measure (with bin width 2Δ*t*). The Jimbo et al. index is normalised by the auto-correlation value at the origin, but the form of this normalisation was not specified. Two versions of multiplicative normalisation were tested, one used the above quantity and the other used its square root. The requirement of the event synchronisation measure (Kreuz et al., 2007b) that Δ*t* should be smaller than the smallest within-train ISI was relaxed so Δ*t* was set freely. The Schreiber et al. (2003) similarity coefficient and the Kruskal et al. (2007) correlation measure had their exponential/Gaussian filters (respectively) replaced with a boxcar filter of width 2Δ*t*. This does not affect whether a measure fulfils the required criteria, but means that a window of synchrony is used to assess correlations (as with the correlation index). A boxcar filter is also more computationally efficient to calculate than Gaussian or exponential filters which require an extra parameter, namely a filter cut-off point, to make computation feasible. Mutual information (Li, 1990) was altered to smooth spikes with a boxcar filter before calculation. The Ripley (1976) *K_mm_* function (a directed measure which measures the correlation of one train *to* another) was made symmetric by setting the correlation measure to be equal to the mean of the two directed variants.

### Evaluating the measures for the necessary and desirable properties

Each measure was tested extensively for the necessary and desirable properties (see Table 2) on a range of synthetic data. This was used instead of experimental data as it is possible to independently alter the key properties (such as rate or correlation). Synthetic data was generated from the following models which replicate four types of experimentally observed spiking patterns: Poisson spiking, Poisson bursting, regular out-of-synchrony spiking and out-of-synchrony bursting. Sample data are presented in Figure 2 and parameter ranges for all four models appear in Table 1.

**Figure 2:**
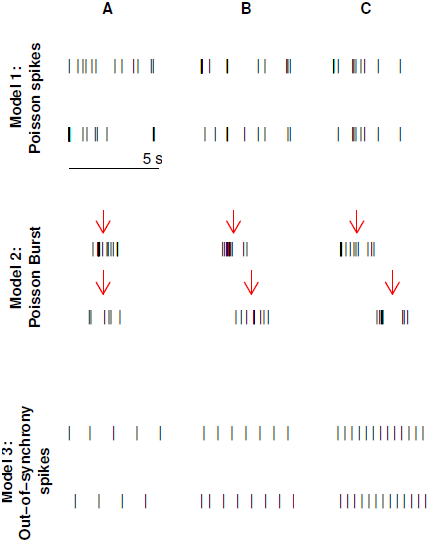
Examples of simulated data used to test measures. Data generated from Model 1 used a Poisson spiking model where both neurons fire at 1.5 Hz with increasing percentage of spike times which are shared with a spike in the other train: 0% (A), 87% (B) and 99% (C). Recording duration *T* = 300 s. Data generated from Model 2 used a Poisson burst model with a burst rate of *λ* = 0.05 Hz, where the number of spikes in each burst is drawn from a Poisson distribution with mean *N* = 8. The positions of the spikes relative to the centre of the burst (indicated by a red arrow) are drawn from a uniform distribution on [1, 1] s (*σ* = 2). The centre of the burst of the second train is offset from the centre of the first by a fixed amount: *O* = 0 s (A), 1 s (B) and 2 s (C), *T* = 3600 s. Data generated from Model 3 shows regular out-of-synchrony firing with increasing firing rate generated using an integrate-and-fire model (see Methods for details). The firing rates are 0.76 Hz (A), 1.27 Hz (B) and 2.5 Hz (C), *T* = 3000 s.

**Table 1:** Parameter ranges used in models to generate test data. **Note that the out-of-synchrony bursting model is phenomenological and so parameters do not related to any particular biological process and are unitless.**

| Parameter | Description | Value |
| --- | --- | --- |
| **Common to all models** |  |  |
| $\Delta t$ | Window of synchrony | (0.01 to 1) s |
| T | Recording Time | (10 to 6000) s |
| **Poisson spiking model** |  |  |
| $\lambda_A, \lambda_B$ | Firing rates of A and B | (0.01 to 5) Hz |
| $\lambda_S$ | Rate of synchronous spikes | (0.01 to 5) Hz |
| **Poisson bursting model** |  |  |
| $\lambda$ | Burst-rate of master train | (0.01 to 0.2) Hz |
| $p$ | Probability of burst in second train | 0 to 1 |
| $O$ | Parameter controlling offset of burst-centre in second train<br>(either fixed or $O$ = range of continuous uniform distribution,<br>or $O$ = variance of Gaussian distribution) | (0 to 2) s |
| $N$ | Parameter controlling number of spikes per burst<br>(either fixed, $N$ = mean of Poisson distribution,<br>or $N$ = Maximum of discrete uniform distribution) | 1 to 20 |
| $\sigma$ | Parameter controlling position of spikes relative to burst centre<br>($\sigma$ = range of continuous uniform distribution,<br>or $\sigma$ = variance of normal distribution) | (0.05 to 2) s |
| **Out-of-synchrony spiking model (Dayan and Abbott, 2001)** |  |  |
| $E_S$ | Synapse reversal potential | (−70 to 0) mV |
| $E_L$ | Resting potential | −70 mV |
| $V_{th}$ | Threshold potential | −54 mV |
| $V_{reset}$ | Reset voltage after action potential | −80 mV |
| $\tau_m$ | Membrane time constant | (0.05 to 1.5) s |
| $r_m \bar{g}_s$ | Specific membrane resistance multiplied by<br>specific membrane conductance | 0.05 |
| $P_{max}$ | Maximum value of synaptic release probability | 0.5 |
| $R_m I_e$ | Change in potential due to injection of small current | 18 mV |
| $\tau_s$ | Synaptic time constant | 0.05 s |
| **Out-of-synchrony bursting model (Shi and Lu, 2009)** |  |  |
| $\alpha$ | Map control parameter | 5 to 30 |
| $\mu$ | Ensures slow gating process is slow | 0.001 |
| $\sigma_A, \sigma_B, \sigma_C$ | Regime-control parameters for neurons A, B and C | −0.4 to −0.1 |
| $g_c$ | Coupling strength | −0.4 to −0.01 |

**Table 2:** Necessary (N) and desirable (D) properties for a correlation measure. Each property is assigned an identifier for ease of reference.

| Necessary properties |  |
| --- | --- |
| N1 | **Symmetry:** The measure $C$ should be symmetric: for spike trains A and B, $C(A, B) = C(B, A)$. |
| N2 | **Robust to variations in the firing rate:** For instance, given two spike trains with a particular correlational structure, if the rates of both trains are doubled but the structure is preserved, the correlation measure should remain the same. |
| N3 | **Robust to amount of data:** In practice this often means robust to recording duration. |
| N4 | **Bounded:** The measure should be bounded taking a value of +1 when the spike trains are identical, with a value of zero corresponding to no correlation and -1 corresponding to anti-correlation. |
| N5 | **Robust to variations in $\Delta t$:** Small variations to $\Delta t$ should not introduce artefacts into the measure. |
| N6 | **Anti-correlation:** The measure should discriminate between no correlation and anti-correlation. |
| Desirable properties |  |
| D1 | **Periods when both neurons are inactive should be ignored:** Periods where both neurons are silent should not be counted as correlated. Experimental data frequently has large periods of quiescence (Wong et al., 1993; Demas et al., 2006) which, if counted would distort calculated correlations. |
| D2 | **Minimal assumptions on structure:** The measure should not assume that spike times have a given underlying distribution as this will lessen the general applicability. |
| D3 | **Minimal Parameters:** The main free parameter in the measure should be the time window of synchrony $\Delta t$. The number of other parameters should be kept to a minimum. |

#### Poisson spiking model

This model assumes that spike trains A and B fire Poisson-distributed spikes with rates *λ_A_* and *λ_B_* respectively. A certain proportion of these spikes are synchronous with a spike in the other train, this forms a Poisson process with rate *λ_S_* . The rates, correlation, recording duration and Δ*t* were varied in isolation to test measures for the required properties.

#### Poisson burst model

The Poisson burst model was used to generate data which replicated the burst-like firing seen in spontaneous retinal waves (a burst of firing when a cell participates in a wave and silence between waves). The model is a doubly stochastic process: the positions (centre-points) of bursts are generated first, and then the bursts themselves (consisting of the number of spikes in the burst, and a position for each spike) are generated.

The first spike train is the “master” train and the centre-points of its bursts are generated according to a Poisson process with a given rate *λ*. The centre-points of the second train are a copy of those in the first train, but with some deleted (each centre-point is deleted with probability *p*). The remaining centre points in the second train are then jittered from their initial positions. This is controlled by a parameter *O* which is either a fixed amount, the variance of a Gaussian distribution or the range of a continuous uniform distribution (in both cases the distributions have zero mean) from which the off-sets are drawn.

For each centre-point of a burst (in either train) the number of spikes in that burst is then generated. This is controlled by a parameter *N* which is either a fixed number or is the mean of a Poisson distribution or the maximum of a discrete uniform distribution from which the number of spikes are drawn. The position of each spike relative to the centre-point is then generated and is controlled by a parameter *σ* which is either the variance of a Gaussian distribution or the range of a continuous uniform distribution (with zero mean in both cases) from which the relative positions are drawn.

The choice of distributions affected all measures consistently and did not qualitatively affect results. The parameters of this model were varied in isolation to test measures for the required properties.

#### Out-of-synchrony spiking model

Regular, out-of-synchrony individual spikes were generated according to a simple inhibitory integrate-and-fire model as described in Dayan and Abbott (2001). Parameters were varied in isolation.

#### Out-of-synchrony bursting model

Out-of-synchrony spike bursts were generated according to a map-based model for neuron membrane voltage (Shi and Lu, 2009). This model was simulated with three neurons and the spikes from one were discarded to produce two spike trains with out-of-synchrony burst-like firing and periods of quiescence. Parameters were varied in isolation.

#### Testing procedure

Measures were tested for necessary and desirable properties in a two-step procedure. Each measure was assigned a unique number at random so that the measure was blindly evaluated. Step one tested all the anonymised measures for the necessary properties using synthetic data. The list of properties appears in Table 2. The measures were tested methodically for each property using data generated from the above models where parameters were varied in isolation (line searches) and the values tested included the experimentally observed ranges (see Table 1 for ranges used). Necessary properties N1–5 were tested using data generated from both the Poisson spiking and Poisson burst model.

As an example, necessary property N3 states that measures must be robust to the recording duration. To test this, data was generated from the Poisson spiking model with rates and Δ*t* fixed, and with varying recording time. Each measure was then calculated. Ten repeats were performed and then one of the fixed parameters was changed and this process was repeated. This was done for several different values of each fixed parameter. This process was then repeated with data from the Poisson burst model. Since the necessary property required that measures are robust to recording duration, measures which showed dependency were judged to lack this property and were not considered further.

After testing for necessary properties N1–5 remaining measures were tested for their ability to distinguish anti-correlation from no correlation (property N6). This property was tested using line searches on data generated from the out-of-synchrony spikes and out-of-synchrony bursts models. Measures which were tested on data from all four models were tested against 7,640 pairs of spike trains in total.

Four measures were shown to possess all necessary properties, which were then assessed on the basis of the desirable properties. Since the desirable properties concern features of the measures (see Table 2), it was not possible to proceed without identifying the measures. At this point the simulation results from the Poisson spiking model were confirmed by analytical calculations for all measures which were tractable under this model.

The second step of testing involved assessing measures for whether they contain extra parameters, whether they count quiet periods as correlated and whether they make assumptions about the statistical properties of the data (Table 2, D1–3). If a measure was shown to lack a desirable property, simulated data from one of the above models was used to show that this had a qualitative effect. These measures were also extensively tested on experimental data to verify that their behaviour on synthetic data was representative of that on experimental data.

## Measures possessing all the necessary properties

To make our article self-contained we briefly present the three previously published measures which possessed all necessary properties.

### Spike count correlation coefficient

The recording time is partitioned into *N* = *T/d* bins of width d (let *d* = Δ*t*, so bin width is equal to the timescale of interest). The spike times **a** and **b** are binned into vectors **A** and **B** of spike counts. The spike count correlation coefficient is Pearson’s correlation coefficient *r* between **A** and **B**:

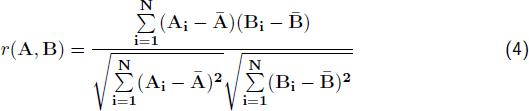

where **Ā** denotes the mean of **A**.

### Kerschensteiner and Wong (2008) correlation

The recording time is partitioned into *N* = *T/d* bins. The spike times **a** and **b** are binned into vectors **A** and **B** of spike counts. A sliding window width *ω* (*≥* Δ*t*) is defined, which is used to calculate local average spike counts. For simplicity let *ω* = (2*n* + 1)Δ*t* for some integer *n*. The Kerschensteiner and Wong correlation, *k* (Equation 5), is Pearson’s correlation coefficient with local average spike counts (**Ã**(**i**)) replacing the global average (**Ā**):

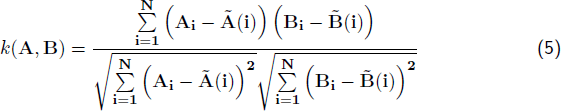

where the local average at bin *i* is given by:

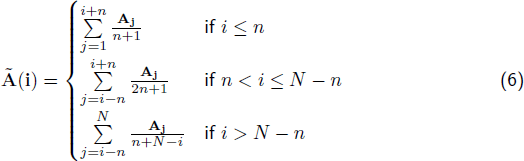

similarly for B**͂**(i).

### Altered Kruskal et al. (2007) correlation

The spike trains are represented as continuous signals, *A* and *B*.

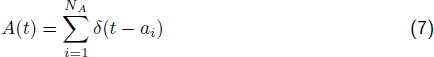

where *δ* represents the Dirac delta function and *B* is represented similarly. These signals are then convolved with a boxcar filter *F* of width 2Δ*t*:

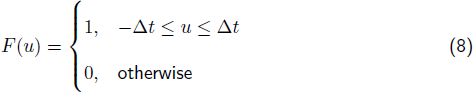

The resulting signal *A*′ is

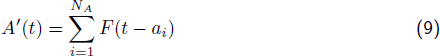

*B*′ is found similarly. The correlation *c* is Pearson’s correlation coefficient of *A*′ and *B*′

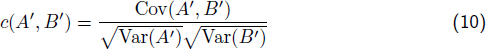

where

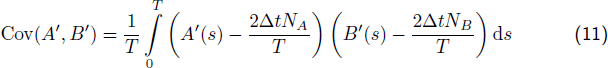

and Var(*A*′) = Cov(*A*′*, A*′). Note that ^(2Δ*tN_A_*)^/*T* is the mean value of signal *A*′ (similarly for signal *B*′).

## Results

### Correlation index

The correlation index is a popular method for quantifying pairwise correlations in neuronal spike times. Given neurons A and B it is defined as the factor by which the firing rate of A increases over its mean value if measured within a fixed window of spikes from B (see Methods). It is the standard correlation measure in studies of spontaneous retinal activity and is also used in several other systems, for instance motor (Personius and Balice-Gordon, 2001) and hippocampal (MacLaren et al., 2011) neurons. It is widely used to compare correlations across different genotypes and ages to infer the function of correlated activity. Since neuronal firing patterns are complex and correlation does not vary in isolation (many other statistics of the data also vary), it is important that measures of correlation are not confounded by other statistics as this means correlations cannot be fairly compared and subsequent inferences are unreliable.

In Methods, we showed that the correlation index is confounded by firing rate, by assuming a neuronal spiking model and calculating an expression for the correlation index which was rate-dependent (Equation 2). To show that this is significant, we use the example of the auto-correlation of a Poisson spike train (the correlation index of this train compared to itself). The correlation index in this case is given by Equation 3 from which it is clear that the rate-dependence is such that the correlations of neurons with low firing rates are up-weighted compared to those with high firing rates. This result was verified by calculating the correlation index of a simulated Poisson neuron compared to itself for varying firing rates (Figure 3). In this case, the correlation index should be constant since no pair of identical spike trains is more correlated than another. In fact, the correlation index decreases with firing rate. The range of firing rates (0.1–5 Hz) used is typical of recordings of spontaneous activity. For example, Demas et al. (2006) reported mean firing rates for four different mouse genotypes at four different ages ranging from 0.45*±*0.04 Hz to 2.15*±*0.22 Hz. The coincidence window Δ*t* was set to 50 ms unless otherwise specified.

**Figure 3:**
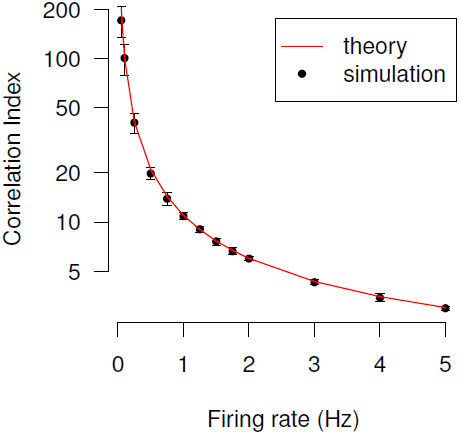
The correlation index is dependent on firing rate. The correlation index of two identical Poisson spike trains is plotted for varying firing rates. Simulation values were generated by simulating one Poisson spike train and then calculating the correlation index comparing this train with itself. Means with error bars of ±1 standard deviation are plotted for ten trials, each of duration 300 s. The theoretical expected value of the correlation index under this model (red line) is given by Equation 3.

A further issue is that the correlation index is unbounded above (Equation 1 and Figure 3). The range of values of positive correlation is [1*, ∞*] whilst that of negative correlation is [0, 1]. Low firing rates return very high values of correlation (see Figure 3). These high values are frequently excluded as outliers, but high correlation index does not imply extreme firing patterns. This makes comparing correlations problematic. For instance, the auto-correlation index of a Poisson neuron with rate 0.1 Hz is nearly twenty times that with rate 1 Hz (Figure 3), however in both cases identical spike trains are being compared, so the conclusion that one is more correlated than the other is erroneous. There is no intuitive feel for how a correlation of 200 compares to a correlation of 10.

### Necessary and desirable properties for a correlation measure

Since the correlation index is confounded by firing rate an alternative measure should be found which is independent of firing rate and can fairly compare correlations. It should be able to replace the correlation index in all analyses. Therefore, since the correlation index is a single-valued measure, the replacement should also be single-valued (as opposed to multi-valued e.g. cross-correlogram). In practice many multi-valued measures can be reduced to single-values by considering just one of their values. The correlation index quantifies correlations over a fixed, small timescale, so its replacement should do the same.

Additionally, there are other properties needed for a measure to fairly compare correlations across recordings where other statistics vary. There are also some desirable properties which either affect the range of correlation values seen in experimental data or require extra information before correlations can be fairly compared. We specified six necessary and three desirable properties, for which we assess potential replacement measures. These are listed in Table 2.

Many measures which quantify the degree of coordination or correlation in neural spike trains exist: an extensive literature search found 33 examples. There is variation in their terminology (such as “coefficients” or “indices”). We use the term “measure” to provide a general term, in the sense that they all “measure” correlations. We classified the measures into six categories:

1. Measures which calculate a distance between spikes trains, or those which calculate a cost involved with transforming one train into another.
2. Measures based on the cross-correlogram, that is, measures counting pairs of spikes which occur within *±*Δ*t* of each other (i.e. the count in the bin centred on zero of the cross-correlogram before normalisation) which is then normalised using some statistic from the cross-correlogram (or justified with reference to it).
3. Measures which also count pairs of spikes which occur within *±*Δ*t* of each other, but which are not derived from the cross-correlogram (e.g. the correlation index).
4. Measures from Information Theory.
5. Measures which consider spike times as a shot-noise process — a term from electronics which considers spike times as discrete events and uses convolution with fixed kernels to derive useful measures.
6. Measures which consider spike times as a marked point process — a concept from Statistics: a point process is a process for which any one realisation consists of a set of isolated points in some space. A marked point process is a point process where additional data exists on the points (other than their location), this data is termed “marks”, in this case a binary “mark” denoting from which neuron the spike originated.

A list of the measures and their classification appears in Table 3. To be as thorough as possible, we have included a broad range of measures, not just those which quantify correlation (some measure synchrony, some similarity and some distance — see Discussion). As a consequence, if a measure is shown to be unsuitable to replace the correlation index, this is no judgement of its usefulness or worth. It is likely that it was designed for use on a different problem and the quantity which it measures is not similar enough to the correlation as we wish to measure it for it to be appropriate in this case.

**Table 3:** Correlation measures evaluated in this study with evidence (if any) for rejecting them as a replacement for correlation index. All measures investigated are arranged according to our devised classification (see Methods). Asterisk denotes that the measure was altered to make it applicable (see Methods). The third column contains an identifier (see Table 2) corresponding to one property which the measure was shown to lack (if any). The fourth column denotes which Figure presents evidence for the lacking property.

| Measure Number | Measure Name | Lacked Property | Evidence in Figure |
| --- | --- | --- | --- |
| Distance measures and Cost functions |  |  |  |
| 1 | Victor and Purpura (1997) | N3 | 5A |
| 2 | ISI-distance (Kreuz et al., 2007a) | N2 | 5B |
| 3 | Hunter-Milton similarity (Hunter and Milton, 2003) | N6 | 5C |
| 4 | Van Rossum (2001) | N3 | 5A |
| 5 | SPIKE (Kreuz et al., 2013) | N2 | 5B |
| Cross-correlation based |  |  |  |
| 6 | Coincidence index (Pasquale et al., 2008) | N2 | 4 |
| 7 | Altered Coincidence index * | N2 | 4 |
| 8 | Cross correlation coefficient (Pasquale et al., 2008) | N2 | 4 |
| 9 | Schreiber et al. (2003) similarity coefficient | N6 | 5C |
| 10 | Altered Schreiber et al. similarity coefficient * | N6 | 5C |
| 11 | Kerschensteiner and Wong (2008) cross-correlation | D3 | 6D |
| 12 | Jimbo and Robinson index (Jimbo et al., 1999) | N2 | 4/5A |
| Synchrony not from Cross-correlation |  |  |  |
| 13 | Correlation index (Wong et al., 1993) | N2 | 3 |
| 14 | Activity pair (Eytan et al., 2004) | N2 | 4 |
| 15 | Unitary events analysis (Grun et al., 2002) | N2 | 4 |
| 16 | Event synchronisation (Kreuz et al., 2007b) * | N2 | 4 |
| 17 | Joris et al. (2006) correlation index | N2 | 4 |
| Information Theory |  |  |  |
| 18 | Mutual information (Li, 1990) | N2 | 4 |
| 19 | Mutual information with smoothing * | N2 | 4 |
| Measures from Shot-Noise Process |  |  |  |
| 20 | Coherence (at zero) (Eggermont, 2010) | N6 | 5C |
| 21 | Spike count correlation (Eggermont, 2010) | D1 | 6A |
| 22 | Smoothed spike count correlation (Kruskal et al., 2007) * | D3 | 6C |
| 23 | Spike count covariance (Eggermont, 2010) | N2 | 4 |
| Measures assuming a Marked Point Process |  |  |  |
| 24 | Stoyan's $K_{mm}$ function (Stoyan and Stoyan, 1994) | N2 | 4 |
| 25 | Isham's mark correlation function (Isham, 1985) | N2 | 4 |
| 26 | Ripley's $K_{mm}$ function (Ripley, 1976) | N2 | 4 |
| 27 | Simpson (1949) index | N2 | 4 |
| 28 | Simpson (1949) index no correction | N2 | 4 |
| 29 | Stoyan's mark covariance function (Stoyan, 1984) | N2 | 4 |
| 30 | Mark variogram (Cressie, 1993) | N2 | 4 |
| 31 | Mark covariance function (Cressie, 1993) | N2 | 4 |
| 32 | Mark conditional expectation (E) (Schlather et al., 2004) | N2 | 5B |
| 33 | Mark conditional variance (V) (Schlather et al., 2004) | N2 | 5 |
| 34 | Mark conditional standard deviation (Schlather et al., 2004) | N2 | 5 |
| Tiling-based |  |  |  |
| 35 | Spike Time Tiling Coefficient | PASS |  |

No measure from the literature was proposed as a replacement for the correlation index and none obviously possesses the full list of necessary and desirable properties. We therefore devised a new measure which conforms to all the criteria — the Spike Time Tiling Coefficient (see Figure 1 for details).

### Step One: evaluating the measures for necessary properties

The replacement for the correlation index must be able to fairly quantify correlations for a wide range of neuronal spiking patterns and therefore possess all necessary and, ideally, all desirable properties. Measures were tested for these properties both analytically and on a wide range of simulated and experimental data. Simulated data is more useful here as individual properties can be altered independently. If a measure lacks at least one necessary property, it was removed from consideration (for brevity, we only present evidence that a measure lacks one necessary property — some lacked more than one). A full list of measures used and its primary reason for rejection (unless it passed step one) are presented in Table 3. Measures were anonymised to remove any possibility of bias. Our procedure was to test for each necessary property in turn (see Methods for details).

From the initial 35 measures, 34 were symmetric (satisfied property N1). The one asymmetric measure: Ripley’s *K_mm_* function was altered to make it symmetric (see Methods).

All 35 measures were tested to ascertain if they were independent of firing rate (property N2). Twenty-five measures showed dependency on firing rate and were therefore rejected. Of these, 22 showed this dependency in the test case of the auto-correlation of a Poisson spike train with varying rate. Since no auto-correlation is more correlated than any other, values should not vary with rate. Any measure which showed rate-dependence was therefore removed according to property N2 (Figure 4). The remaining three measures which lacked property N2 were independent of rate for auto-correlation, but showed rate-dependency when firing rates differ. This dependency is clear in the test case of two independent Poisson spike trains, one with fixed rate and the other with varying rate. Measures whose correlation varied with the rate of the second train were removed (Figure 5A). Note that two versions of measure 12 were considered (see Methods); both lacked property N2 (one version appears in Figure 4 and the other in 5A). For counting purposes, we consider Measure 12 as one measure which lacked property N2 as evidenced by considering auto-correlations. Typically, measures are confounded by firing rate because it is used in their normalisation.

**Figure 4:**
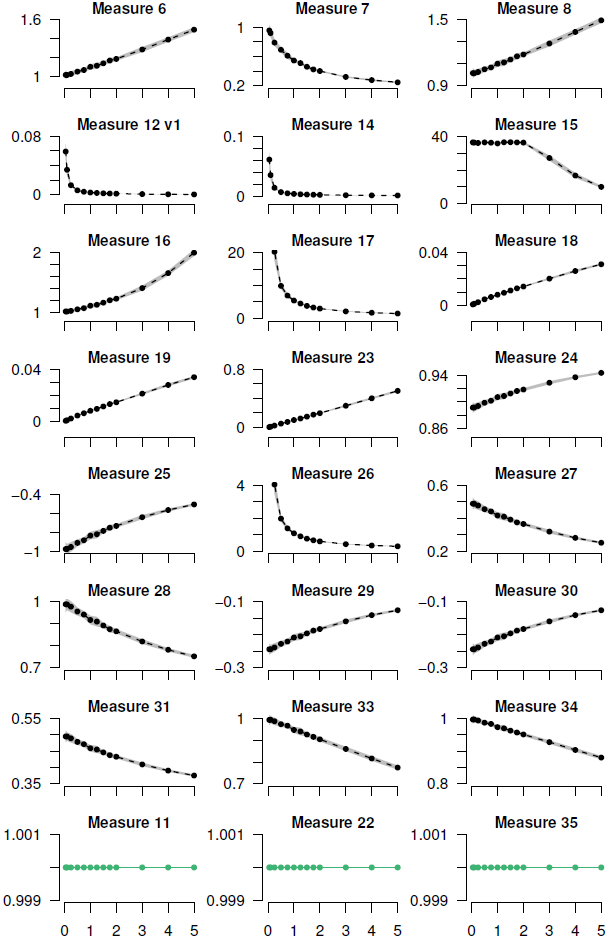
Twenty-one measures are rejected since they are dependent on firing rate (lack property N2). All measures which showed rate-dependence when tested on the auto-correlation of Poisson spike trains are plotted. Three measures which did not show rate-dependence are also shown in the bottom row for comparison (green). One Poisson spike train was simulated for 300 s for varying rates (0.05–5 Hz) and the measures were calculated comparing this spike train to itself. Means of ten repeats are plotted and ±1 standard deviation is shown by grey shading. The identity of each measure appears in Table 3. Note that the correlation index is not presented here, but in Figure 3 and that measure 12 has two versions; one appears here and the other in Figure 5A, see Methods for details.

**Figure 5:**
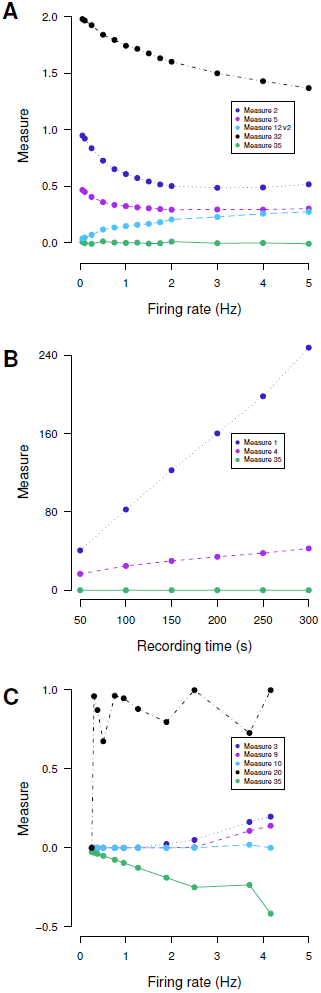
Nine measures are rejected using remaining necessary properties. A: Measures which are dependent on firing rate (lack property N2) where dependency is not obvious from auto-correlation are applied to data generated from the following Poisson spiking model: one train has rate 3 Hz and the other’s rate varies (0.1–5 Hz). There are no shared spike times (*T* = 300 s). The second version of Measure 12 is shown (the first version is in Figure 4), see Methods for details. B: Measures which are dependent on recording time (lack property N3) are applied to data generated from the following Poisson spiking model: both neurons fire at rate 1 Hz and 10% of spike times are shared. The recording duration varies from 50–300 s and Δ*t* = 0.6 s (higher than usual since for these measures smaller values cause issues with computational precision). C: Measures which cannot distinguish anti-correlation from no correlation (lack property N6) are applied to regular out-of-synchrony spikes of varying rate (0.25–4.2 Hz) generated using an integrate-and-fire model described in Dayan and Abbott (2001), Figure 5.20. The parameters are as in their figure with the following exceptions: *P_max_* = 0.5, *R_m_I_e_* = 18 mV. *τ_s_* varied from 0.05–1.5 s and *E_s_* from 0 mV to 70 mV, *T* = 3000 s. In all panels, Measure 35 (which possesses the necessary properties) is shown for comparison (green), means of ten repeats are plotted and error bars are omitted for visual clarity.

The remaining ten measures were tested to ascertain if they were robust to the amount of data available (property N3) which in practice is proportional to the recording time. Measures should therefore be independent of this (within experimental ranges — minimum two minutes with usual range 20–100 minutes (Eglen et al., 2014)). To test this, two Poisson spike trains with fixed rates and correlations were simulated for different recording times. Two measures showed dependency and so were rejected (Figure 5B). These measures were a distance measure and a cost function which are not normalised and so increase as the number of spikes increases.

The eight remaining measures were then tested for the correct range (property N4). They should be equal to +1 for identical spike trains, 0 for no correlation and -1 for strong anti-correlation. All measures were bounded and therefore could be scaled to have the required range provided that they can discriminate between no correlation and anti-correlation (property N6) and so none were rejected at this point.

All eight measures were found to be robust to small variations in Δ*t* (property N5). Of these eight measures, four were rejected since they could not distinguish no correlation from anti-correlation (property N6). The test case on which these measures were removed was regular out-of-synchrony spiking (Figure 5C). The measures rejected were measures of similarity, rather than correlation which therefore could not distinguish this firing pattern from independent spike times.

In summary, 31 out of 35 measures were rejected in step one as they lacked at least one necessary property. A list of each measure considered and the reason for its rejection can be found in Table 3.

### Step Two: selecting one measure based on desirable properties

The four measures possessing all necessary properties were (1) the spike count correlation (Eggermont, 2010) (2) the Kerschensteiner and Wong (2008) measure (3) an altered version of the correlation measure from Kruskal et al. (2007) and (4) the Spike Time Tiling Coefficient. A brief description of each measure is given in Methods. Selecting one measure proceeded on the basis of desirable properties (Table 2). These affect either the range of correlations (D1: periods of silence should not count as correlated), the applicability of the measure (D2: spike times should not be assumed to follow a particular distribution), or mean that extra information is required to compare correlations (D3: extra parameters are discouraged).

The spike count correlation calculates the Pearson’s correlation coefficient of binned spike counts. We set our bin width to Δ*t* (see Methods). Since firing rates are sparse, a large proportion of bins have zero spikes in both trains, which counts as correlated. This distorts the values of correlations making results difficult to interpret (Figure 6A and B). The spike count correlation therefore lacks property D1 but does possess properties D2 and D3. A further limitation is that spikes which are within *±*Δ*t* of each other may fall into different bins and these coincidences are missed. It has also been reported that the spike count correlation increases with firing rate (De la Rocha et al., 2007) which lessens its ability to compare correlations fairly. This is not reported here since variation in firing rate was small compared to variation across trials on the time scales considered.

**Figure 6:**
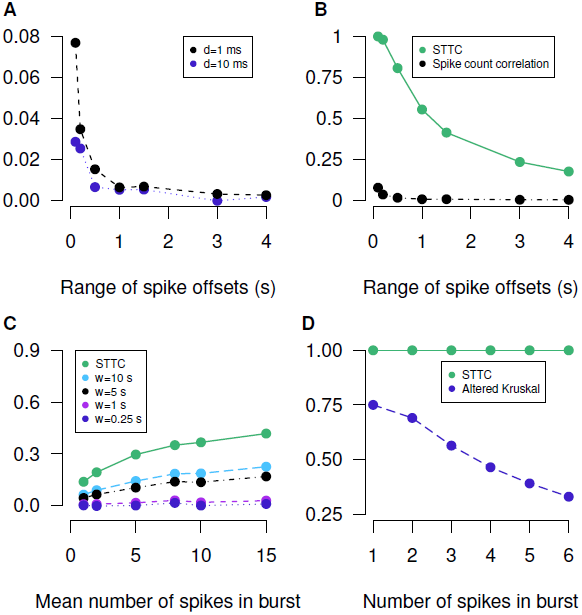
Detailed examination of the four measures which possess all necessary properties eliminates three on the basis of the desirable properties. A: The spike count correlation with different bin widths (d — see Methods) is applied to data from the following Poisson burst model which has increasing range of spike offsets within a burst: both neurons have a burst rate of 0.05 Hz, burst centres have 0 s offset, each burst contains 8 spikes whose positions are drawn from a uniform distribution of varying width (0.1–4 s) centred on the burst centre (*T* = 3600 s). B: The Spike Time Tiling Coefficient (STTC) applied to identical data to that in A. The spike count correlation with *d* = 1 ms is plotted (black) for comparison. C: The Kerschensteiner and Wong correlation measure with different lengths of the averaging window (*w* — see Methods) is applied to data from the following Poisson burst model with increasing number of spikes per burst: both neurons have a burst rate of 0.05 Hz, burst centres have 0 s offset, the number of spikes in each burst is drawn from a Poisson distribution with increasing mean (from 1 to 15). Spike positions are drawn from a uniform distribution of width 2 s centred on the centre of the burst (*T* = 3600 s). D: The Kruskal measure is applied to spike times from the following model: two independent Poisson neurons are simulated each with rate 0.1 Hz. For each spike (in either train) a burst is generated in the other train with 0 s offset of the burst centre and 1–6 spikes whose positions are drawn from a uniform distribution of width 2Δ*t* around the burst centre (*T* = 2000 s, Δ*t* = 0.1 s). The Spike Time Tiling Coefficient (green) is plotted in panels C and D for comparison. For all panels the mean of ten repeats is plotted, error bars are omitted for visual clarity.

The Kerschensteiner and Wong measure is an altered version of the spike count correlation which replaces the global average firing rate with a local average firing rate (using a sliding window) which prevents periods of silence counting as correlated. Data is binned into bins of width Δ*t* and the sliding window calculates a local average across a fixed number of bins. Whilst the alteration is effective, this measure possesses desirable properties D1 and D2, it introduces an extra parameter: the length of the sliding window (discouraged by property D3). In order for this measure to be informative, the length of the sliding window must be optimised for the data. If the sliding window is too large compared to the periods of quiescence, then when both trains are quiet, the correlation at that point will be positive. If it is shorter than the burst lengths, then periods where one neuron has a burst and the other does not will count as zero (should count as anti-correlated) since in the silent train both the local average and the individual spike counts will be zero (see Equation 5). Therefore the measure varies qualitatively with this parameter (Figure 6C), especially on burst-like data. Correlations cannot be fairly compared if this parameter varies, which it is likely to since a poor choice of parameter can lead to uninformative values of correlation.

The altered version of the Kruskal et al. measure changes the spike count correlation to overcome the fact that coincidences may be missed if they fall in different bins: it smooths spike trains with a boxcar kernel before calculations. This does not possess property D1 as silence is still counted as correlated. It does possess property D2 and D3 (since it is possible to calculate exactly).

The Spike Time Tiling Coefficient does not count periods of quiescence as correlated and thus possesses property D1. It does not assume a statistical distribution of spike times and therefore possess property D2. The only free parameter is Δ*t* and it therefore possess property D3.

Whilst the Kruskal et al. measure lacks necessary property D1 which reduces the range of correlations produced, this effect is not as large as for the spike count correlation. We also note that the Kruskal et al. measure is like a “similarity measure” in the sense that it takes value +1 only if the spike trains are identical whereas the Spike Time Tiling coefficient is equal to +1 for a wider range of firing patterns (those where *P_A_* = *P_B_* = 1) and identical trains can be distinguished from those which are merely highly correlated by letting Δ*t →* 0. Since the Kruskal et al. measure is close to a “similarity” measure, relatively low values of correlation can be assigned to highly correlated firing patterns. For instance, consider two spike trains, where if one has a single spike, the other fires several spikes within *±*Δ*t* of that spike and vice versa. This is clearly highly correlated (a firing pattern which is indicative of some relationship between the neurons) although the spike trains are not similar so the Kruskal measure assigns low values of correlation. This value depends on the number of spikes fired when there is one spike in the other train (Figure 6D) and misrepresents the correlation. The Spike Time Tiling Coefficient assigns a correlation value of +1 independent of the number of spikes.

In practice both the Kruskal et al. and the Spike Time Tiling Coefficient were found to be adequate reporters of correlation on experimental data. Since the Kruskal measure lacks property D1 and assigns high correlations only to similar spike trains, the Spike Time Tiling Coefficient is able to pick up a larger range of correlated firing patterns and we therefore recommend it to replace the correlation index.

### Re-analysis of experimental data using the Spike Time Tiling Coefficient

The issues with the correlation index raise questions about the reliability of studies which have used it to draw their conclusions. Since the correlation index is confounded by firing rate, it should not be used to compare correlations in data where rates differ significantly. Firing rates frequently vary across age, phenotype and presence of pharmacological agonists and so this calls into question the results of many correlation analyses: some conclusions about differences in correlations may be due to the confounding effect of the firing rates, and not the correlations themselves. Since the correlation index has been widely used in the field (we found 43 papers which used it, 29 of which have been published since 2008), we also considered the wider implications of its use. In particular, what conclusions were at risk of changing if the data were to be re-analysed using the Spike Time Tiling Coefficient? Four examples using seven studies in the developing retina are presented to demonstrate the use of the Spike Time Tiling Coefficient in place of the correlation index. For this work, we used the freely available retinal wave data from the CARMEN portal (Eglen et al., 2014) (https://portal.carmen.org.uk/).

### Example one: connexin isoforms

Re-analysis of existing recordings of spontaneous retinal waves using the Spike Time Tiling Coefficient in place of the correlation index can show significant differences. As an example, we re-analysed MEA recordings from Blankenship et al. (2011) which compared the statistical properties of wild type spontaneous retinal activity to those from mutant mice lacking either one or two connexin isoforms (Cx45 and Cx36/Cx45). This study reported that the two mutants exhibit substantially higher firing rates compared to wild type (Figure 7A) and when the correlation indices are calculated pairwise and plotted against electrode separation (the standard analysis) the size of the correlation depends inversely on the mean firing rate (Figure 7B). That is, wild type has both the lowest firing rate and the highest correlation and Cx36/Cx45dko has the highest firing rate and the lowest correlation. From the raster plots (Figure 7A) all phenotypes exhibit some correlated firing and the differences in correlation patterns are not as large as the differences in firing rate.

**Figure 7:**
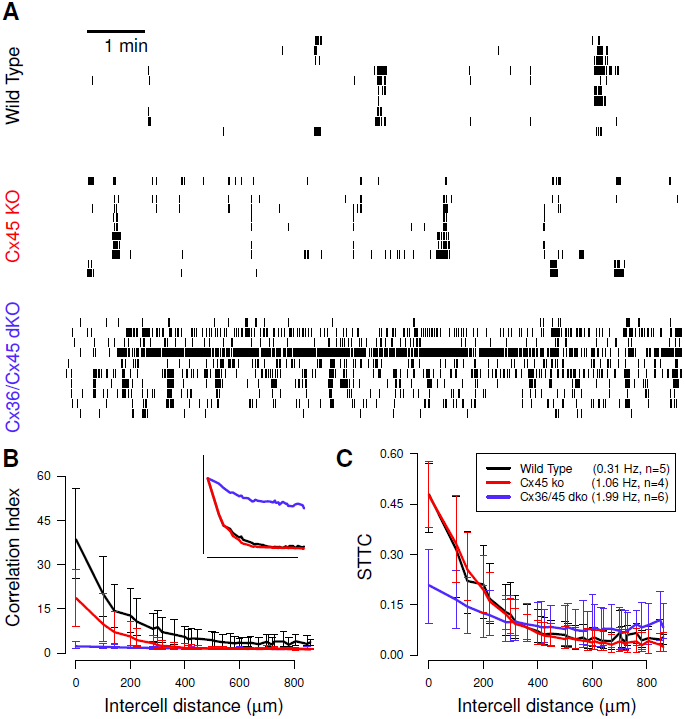
Evaluating correlations in retinal waves recorded from connexin mutant mice shows that the Spike Time Tiling Coefficient can significantly alter conclusions. A: raster plots of ten spike trains over a ten minute interval, recorded from retinas isolated from P12 wild type mouse and two mutant mice (lacking either one or two connexin isoforms-Cx45 and Cx36/Cx45), P11 Cx45ko and P10 Cx36/Cx45dko. Data is from Blankenship et al. (2011) and raster plots follow the presentation of their Figure 2A. The mean firing rate and number of animals (n) from each genotype is recorded in the legend. B: Pairwise correlation index as a function of intercellular distance for each genotype. Data points are medians over all recordings and error bars indicate the interquartile range (IQR). Inset shows the same data normalised (multiplicatively) by genotype so that the correlation indices are identical at zero distance, following Figure 2B in the original publication. C: same as panel B, using the Spike Time Tiling Coefficient (STTC) in place of the correlation index. Compare with both B and B insert. In both B and C Δ*t* = 100 ms as in the original publication. The distances at which correlations are measured are the discrete set of separations possible on the MEA grid.

When the data is re-analysed using the Spike Time Tiling Coefficient, the results are strikingly different (Figure 7C): wild type and Cx45ko have highly similar correlation values and the difference between correlations in wild type and Cx36/Cx45dko are much smaller. Whilst correlations are difficult to judge from the raster plots, those of wild type and Cx45ko have several correlational features in common: they both show waves and also some correlated spiking outside of waves. Waves cannot clearly be seen in the raster plot of Cx36/45dko, although there is correlated firing. The relative correlations of these phenotypes measured using the correlation index reflect the differences in firing rate, whereas, when measured using the Spike Time Tiling Coefficient, they reflect differences in correlational structure (Figure 7A).

In the original publication, the authors noted that the correlation values for Cx36/Cx45dko were so low that it was difficult to deduce anything about its distance dependence relative to the other phenotypes. To make this comparison they (multiplicatively) normalised the correlation indices so that all phenotypes had the same value (the wild type value) at zero distance. From this they deduced that the distance dependence of wild type and Cx45ko were very similar and also that the correlations of Cx36/Cx45dko had a weaker distance dependence than the other two phenotypes. This result is immediately apparent using the Spike Time Tiling Coefficient, without the need to normalise: wild type and Cx45ko have very similar values at all distances and Cx36/Cx45dko has weaker distance dependence. This phenotype has less correlation than the other two at smaller distances and more correlation at greater distances (*≥* 400 µm). This is not apparent using the correlation index; the Spike Time Tiling Coefficient thus provides a more informative comparison of the correlations.

### Example two: developmental changes in correlation

A key result used to support the hypothesis that correlated activity plays a role in map formation is that correlations in spontaneous retinal activity decrease with age both in ferret (Wong et al., 1993) and mouse (Demas et al., 2003). These studies reported different variation in firing rates with age: firing rates decrease with age in ferret, but increase with age in mouse (confirmed by Maccione et al. (2014)). Since neurons with high firing rates have down-weighted correlation indices, it is possible that this result is due to the confounding effect of the increasing firing rates in mouse. This is unlikely to be the case in ferret where firing rates decrease. Re-analysis with the Spike Time Tiling Coefficient confirmed that in both cases, correlations do decrease with age and so this conclusion stands (Figure 8).

**Figure 8:**
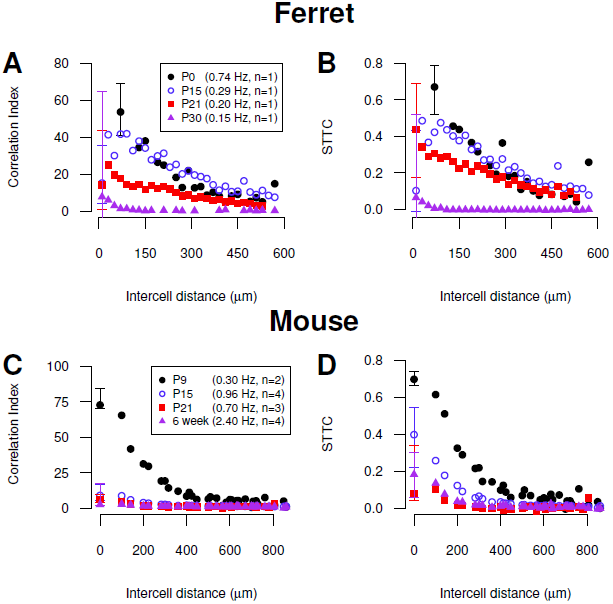
**Re-analysis using the Spike Time Tiling Coefficient supports the conclusion that correlations in spontaneous activity in the developing ferret and mouse retina decreases with age**. A: The correlation index is calculated pairwise and shown as a function of electrode separation for spontaneous retinal activity in developing ferret for four different ages (data from Wong et al. (1993)). The distances at which correlations are measured were binned (bin width 20 µm) due to high density. B: same as panel A using the Spike Time Tiling Coefficient (STTC) in place of the correlation index. C: The correlation index is calculated pairwise and shown as a function of electrode separation for spontaneous retinal activity in developing mouse for four different ages (data from Demas et al. (2003)). The distances at which correlations are measured are the discrete set of separations possible on the MEA grid. D: same as panel C using the Spike Time Tiling Coefficient. In all panels median values are plotted and IQRs are only shown at the smallest separation distance. Other IQRs are omitted for visual clarity. Mean firing rates and number of animals (n) for each age are recorded in the legend.

### Example three: *β*2 genotypes

A key group of genetically modified mice which provide (somewhat controversial) evidence that correlations are key to map formation is the *β*2(KO) mutants (animals lacking the *β*2 subunit of the nicotinic acetylcholine receptor) which form a defective retinotopic map. This mutant was initially thought to have uncorrelated activity (McLaughlin et al., 2003), but it was later shown that retinal waves exist (Sun et al., 2008; Stafford et al., 2009). The *β*2(KO) mice have slightly higher firing rates and weaker distance-dependence of correlations than wild type (typically at short distances these mutants have lower correlation than wild type, but they have higher correlation at large distances). Re-analysis of data from Sun et al. (2008) and Stafford et al. (2009) using the Spike Time Tiling Coefficient confirms reported results (Figure 9).

**Figure 9:**
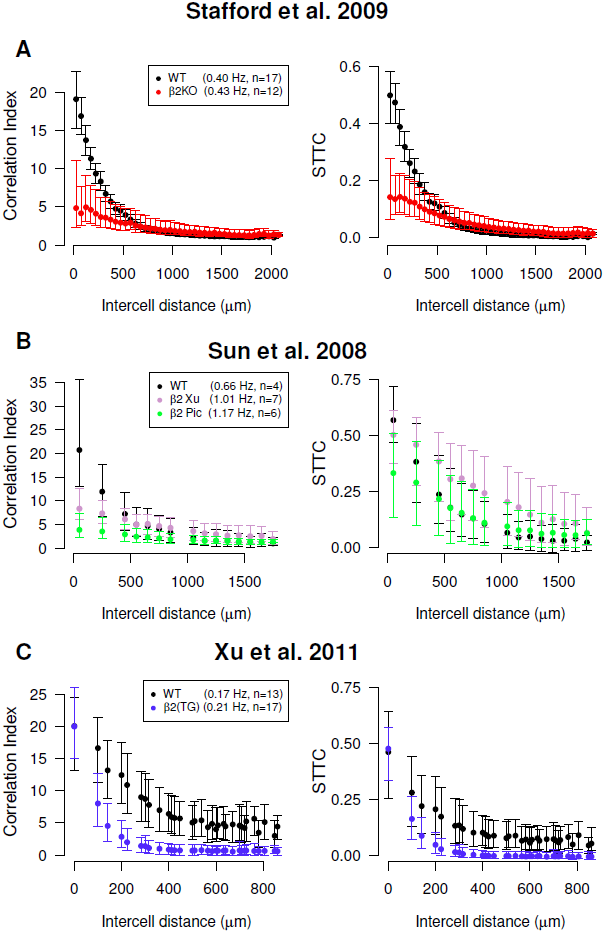
Re-analysis of experimental data using the Spike Time Tiling Coefficient supports the conclusions that the *β*2(KO) and *β*2(TG) mouse phenotypes show lower correlations in spontaneous retinal activity than those of wild type. A: The correlation index (left) or Spike Time Tiling Coefficient (STTC) (right) is plotted pairwise against electrode separation for recordings of spontaneous retinal activity for P6 wild type and *β*2(KO) phenotypes (data from Stafford et al. (2009)). B: The correlation index (left) or Spike Time Tiling Coefficient (right) is plotted pairwise against electrode separation for recordings of spontaneous retinal activity for P5 wild type and two *β*2(KO) phenotypes: Xu and Picciotto (Pic) (data from Sun et al. (2008)). C: The correlation index (left) or Spike Time Tiling Coefficient (right) is plotted pairwise against electrode separation for recordings of spontaneous retinal activity for P4 wild type and *β*2(TG) phenotypes (data from Xu et al. (2011)). In all panels Δ*t* = 100 ms, as in original publications, medians are plotted and the error bars show the IQR. Mean firing rates and number of animals (n) for each phenotype are recorded in the legend. All recordings at 37*^◦^*C. The distances at which correlations are measured are the discrete set of separations possible on the MEA grid.

Although the conclusion stands, reanalysis of the data from Sun et al. (2008) shows differences from the original analysis (Figure 9B): the correlation indices are noticeably confounded by the firing rates which differ across the genotypes. Re-analysis using the Spike Time Tiling Coefficient shows that the differences in correlation between phenotypes are smaller than previously reported; all three phenotypes now show significant correlation at short distances. The order of phenotypes by correlation at short distance (wild type is highest, *β*2(KO) (Picciotto) is lowest) is preserved as is the order by distance dependence (wild type is strongest, *β*2(KO) (Picciotto) is weakest).

Re-analysis of recordings of the *β*2(TG) mouse (Xu et al., 2011) confirm that this phenotype shows weaker correlations than that of wild type (Figure 9).

### Example four: Age-related changes in *β*2(KO)

Retinal waves of *β*2(KO) mutants at different ages (P4–7 and wild type P5–6) were recorded by Kirkby and Feller (2013) as controls in their study of intrinsically photosensitive retinal ganglion cells. Sample raster plots for P4 *β*2(KO) and P5 wild type are shown in Figure 10A. Firing rates of the *β*2(KO) mutants increase (see Figure 10 legend) and their correlation indices decrease (Figure 10B) with age. In addition, P4 and P5 *β*2(KO) are extremely and erratically correlated whereas P6–7 *β*2(KO) and wild type show typical correlations and distance dependence. One might, therefore, conclude that the recordings of P4–5 *β*2(KO) show highly correlated and extremely unusual firing patterns. However, this is not borne out by inspection of the raster plots (see Figure 10A) - the firing patterns of the P4 *β*2(KO) mutant are not too dissimilar to those of wild type, albeit occurring at a much lower rate. We note that the mean firing rates *β*2(KO) P4–5 are low compared to other ages (and wild type) and so this behaviour is likely to be due to the confounding effect of the firing rates. Re-analysis of this data using the Spike Time Tiling Coefficient confirms this (Figure 10C): all correlations and distance dependencies are now within typical ranges. These two phenotypes (*β*2(KO) P4–5) still show more variability in their distance dependencies, but we note that this is likely to be due to the small number of recordings, and that this variability is much less than that seen using the correlation index.

This re-analysis alters the conclusion that correlations in *β*2(KO) genotypes decrease with age (in P4–7), which is caused by the confounding effect of increasing firing rates. Analysis with the Spike Time Tiling Coefficient shows that they tend to increase with age (the order of correlations from lowest to highest is P5,P4,P6,P7).

**Figure 10:**
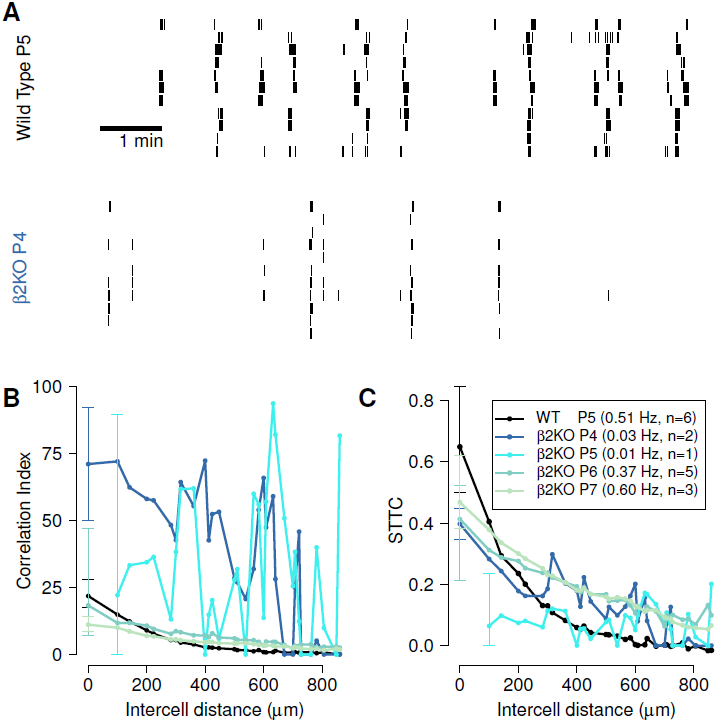
Re-analysis of age related changes in *β*2(KO) mutants shows that the Spike Time Tiling Coefficient is able to more accurately quantify correlations which the correlation index ascribes as extremely correlated. A: Raster plots of ten spike trains over a ten minute interval, recorded from retinas isolated from P5 wild type mouse and P4 *β*2(KO) mouse. All data are controls from Kirkby and Feller (2013). B: Pairwise correlation index as a function of intercellular distance for P5 wild type mouse and *β*2(KO) mouse of different ages (P4–7). C: Same as panel B but using Spike Time Tiling Coefficient (STTC). The mean firing rate and number of animals (n) from each genotype is recorded in the legend. Data points are medians over all recordings and error bars indicate the IQR. For visual clarity, IQRs are only shown at the smallest separation distance. The distances at which correlations are measured are the discrete set of separations possible on the MEA grid. Δ*t* = 100 ms, as in original publication and recordings were performed at 33 *−* 35*^◦^*C.

### Variation of window of synchrony Δ*t* can reveal timescales of correlation

The coincidence window Δ*t* is a free parameter which should be fixed in order to compare correlations. It can also be used to find time scales of correlation in a data set. Figure 11 shows how varying Δ*t* changes the Spike Time Tiling Coefficient (for the data from Example Three). Useful timescales can be found by considering local maxima and minima and the gradient of Spike Time Tiling Coefficient (note that the limit of the Spike Time Tiling Coefficient as Δ*t → T* is +1). For instance in all panels in Figure 11 there is a clear change in gradient around 0.5–1 s which could indicate a timescale of correlation. In panels A and B, the wild type gradient is largest between 0.01 s and 0.05 s as is *β*2(TG) in panel C, which could also indicate a useful scale. Differences in correlational time scales between phenotypes are also apparent, for instance in panel C wild type spikes are less correlated than those of *β*2(TG) on timescales of Δ*t* ≤ 0.1 s and more correlated than *β*2(TG) on larger timescales (note, however, significant overlap of error bars).

**Figure 11:**
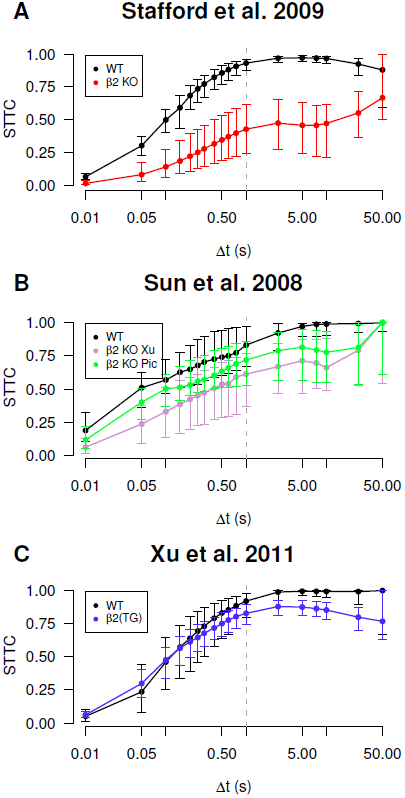
Varying the window of synchrony Δ*t* can be informative about correlational timescales inherent in data. A: The Spike Time Tiling Coefficient (STTC) of spike trains from Stafford et al. (2009) was calculated pairwise (as in Figure 9B) and the median value at the smallest electrode separation is plotted for varying Δ*t*. Error bars show the IQR. The genotypes shown are P6 wild type and *β*2(KO). B: same as panel A, but data from Sun et al. (2008) (see Figure 9D). The genotypes shown are P5 wild type and two *β*2(KO) phenotypes: Xu and Picciotto (Pic). C: same as panel A, but data from Xu et al. (2011) (see Figure 9F). The genotypes shown are P4 wild type and *β*2(TG) mouse. Vertical lines at Δ*t* = 1 s indicate separation between region with strong Δ*t* dependency (Δ*t* ≤ 1) and weaker dependency (note x-axis has a log-scale and that the limit of the Spike Time Tiling Coefficient as Δ*t* tends to infinity is one).

## Discussion

We have described the need to correctly quantify neuronal correlations and considered a popular measure, the correlation index. We have shown that it is confounded by firing rate and is unbounded above which means it cannot fairly compare correlations when firing rates differ significantly. We aimed to find a measure which could be used as a replacement for the correlation index and which could fairly compare correlations. We listed necessary and desirable properties which such a measure needs and found 33 other existing measures of correlation. Since no measure obviously possessed all listed properties, we proposed a novel measure of correlation: the Spike Time Tiling Coefficient. We blindly tested all measures for the properties using synthetic and experimental data. We reiterate that no existing measure was designed to replace the correlation index so the exclusion of a measure is no reflection of its usefulness. Four measures possessed all the required properties and the Spike Time Tiling Coefficient was chosen as the most appropriate replacement on the basis of the desirable properties. To demonstrate its use we re-analysed data from seven studies and showed that it can significantly alter conclusions.

### The form and use of the Spike Time Tiling Coefficient

Both the correlation index and the Spike Time Tiling Coefficient use a small time window to identify spike pairs which are indicative of an overall correlation between the spike trains. We wish to quantify correlations that are functionally significant, which are typically those which can affect synaptic change. In spontaneous retinal waves, correlated activity is thought to contribute to map formation by helping neighbouring neurons wire to common targets via a Hebbian mechanism (Demas et al., 2003). This process has a critical time window: studies of spike-time-dependent-plasticity (Zhang et al., 1998) provide a means to estimate its width, which is age-and species-dependent, around 50–500 ms (Lee et al., 2002). More recently, other rules, such as burst-time-dependent-plasticity, suggest longer windows (Butts et al., 2007).

The time window, Δ*t*, can take any value of interest; often that value is dictated by the phenomenon being investigated. For instance, in spontaneous retinal activity, Δ*t* is dictated by spike-time-dependent-plasticity and in cortical circuits, local oscillatory events could be used to find a Δ*t* of interest as they are reporters of synchrony (Harris et al., 2003). If there is no prior Δ*t* of interest, its value could be varied to show timescales of correlations (see Figure 11). If varying Δ*t* is infeasible, an approximate value of interest could be generated by inspection of cross-correlograms.

The Spike Time Tiling Coefficient assumes stationary spiking patterns. Correlations calculated from highly non-stationary data may be misleading as network states greatly influence firing patterns in e.g. hippocampal firing, so the average value of correlation may not accurately represent the data. Changes in correlation over a non-stationary recording can be identified using the Spike Time Tiling Coefficient by calculating it within a sliding window to get a temporally varying correlation. This window must be large enough to capture representative behaviour and so if it is required to capture changes in correlation on a very small timescale, a measure which incorporates some form of localised measurement (Kerschensteiner and Wong, 2008) may be preferable.

We have used the word “correlation” throughout but noted that the terminology varies. Few of the measures measure correlation in the statistical sense (the degree to which measurements on the same group of elements tend to vary together). Neither the correlation index nor the Spike Time Tiling Coefficient are correlations under this definition. The only “true” correlation is the spike count correlation coefficient (and the altered version).

We suggest that the correct terminology for what we wish to measure is “affinity” in the biological sense — meaning a relationship or resemblance in structure that suggests a common origin or purpose. We wish to measure a relationship between spike times that may indicate that neurons are involved in the same process. In spontaneous retinal activity, we wish to measure the propensity of two neurons to fire close in time to each other in such a way that it can affect their wiring onto a common target.

Pairwise measures of correlation will only capture a subset of the full correlational relationships in a neuronal population. Population dynamics are less noisy than pairwise dynamics and may encode critical information. Methods exist to study higher-order correlations (Nakahara and Amari, 2002; Walters et al., 2008) and investigating population dynamics is a common approach (Okun et al., 2012). We have focused on pairwise correlations, partly due to its popularity and the large literature concerning its quantification but also as evidence suggests that pairwise interactions can account for much of the observed higher-order interactions (Schlens et al., 2006; Schneidman et al., 2006).

### The Spike Time Tiling Coefficient in the re-analysis of data and cross-study comparisons

We have re-analysed the results of seven studies using the Spike Time Tiling Coefficient instead of the correlation index. This has shown that sometimes the correlation index reflects differences in firing rate more than differences in correlation and so the Spike Time Tiling Coefficient can change the conclusions of studies. For example, our re-analysis of the data from Blankenship et al. (2011) significantly changes the conclusions about the relative correlations in spontaneous retinal activity of connexin knock-out mutants.

The popularity of the correlation index means that many studies have drawn conclusions on the basis of a problematic measure. Studies where firing rates differ significantly and the order of phenotypes by correlation is the reverse of the order by firing rate are likely to change. However, of these, we have shown that two important conclusions made using the correlation index still stand when re-analysed with the Spike Time Tiling Coefficient. These are the age-dependent changes in wild type correlations in mouse and ferret (Figure 8) and also relative correlations of wild type and *β*2(KO) mouse mutants (Figure 9).

Although the re-analysis of data from *β*2 mutants broadly confirms the conclusions of the original studies, the correlations of the two *β*2(KO) mutants from Sun et al. (2008) show stronger distance-dependence and values closer to wild type than was evident using the correlation index (Figure 9B). Both mutants have relatively high firing rates (see Figure 9 legend) so their correlation indices are down-weighted, making the distance dependence appear weaker.

The *β*2(KO) mutant line used in Stafford et al. (2009) is the same as the Xu knock-out line used in Sun et al. (2008) with ages P6 and P5 respectively. We note variation between studies of the size and distance dependence of these correlations. Some of the variation between reported correlations may be due to different bath solutions (Stafford et al., 2009), or possibly to age-related differences. However, we also note large cross-study variation in the wild type control (P4 (Xu et al., 2011), P5 (Sun et al., 2008) and P6 (Stafford et al., 2009)). Given that they are similar ages, we would expect control firing rates and correlations to be reasonably similar (Wong et al., 1993), however both the firing rates and maximal correlation values vary significantly (Figure 9). Variation between studies for the controls is large so the differences observed in *β*2(KO) between studies are not surprising given this and the use of different bath solutions.

One conclusion about the correlations in *β*2(KO) which changed when re-analysed using the Spike Time Tiling Coefficient is that correlations tend to increase during P4-7, rather than decrease (see Figure 10). This case provides a good example of how the correlation index can assign extreme values to data which is not atypical but which has low firing rates. These data are typically excluded as outliers: many studies filter neurons for extremely low or high firing rates before calculating correlations (Maccione et al., 2014). However, re-analysis using the Spike Time Tiling Coefficient shows that this is not necessary: meaningful conclusions about correlations can still be obtained from this data.

## Conclusions

Here, we have used spontaneous retinal activity as a case-study. Since quantifying correlations in spike times is of wider interest, we expect the Spike Time Tiling Coefficient to have applications to measuring correlations in other systems, such as hippocampal cultures (Godfrey and Eglen, 2009), multi-sensory integration (Parise et al., 2013) or motor control (Lee and Lisberger, 2013). With regards to our case-study, we hope that its use will help clarify the exact role of correlations in map formation.

## Acknowledgements

The authors thank Keith Godfrey for initial investigations into the limitations of the correlation index that led to this study. We thank Ellese Cotterill, David Dupret, Diana Hall, Mattias Henning, Johannes Hjorth, Evelyne Sernagor and Álvaro Tejero-Cantero, for comments on the manuscript and also Kate Belger for administrative support. The authors thank the Wellcome Trust (SJE; grant number 083205) and EPSRC (CSC) for funding.

## List of abbreviations and notation used

ISI: Interspike interval
IQR: Interquartile range
MEA: Multielectrode array
T: Recording time
Δ*t*: Time window of synchrony
a: Vector of spike times of neuron A
*^N^A*: Number of spikes from neuron A in recording
*N*_*A,B*[−Δ*t*,Δ*t*]_: Number of spike pairs where a spike from neuron A occurs within ±Δ*t* of a spike from neuron B
*^λ^A*: firing rate of spike train A
*^λ^S*: firing rate of spikes shared between two trains
*d*: bin width
A: vector of binned spike counts of A
*ω*: sliding window width
Ā: Global average of **A**
Ã: Local average of **A**
*A*: spike train of neuron A represented as a signal
*F*: convolution kernel
*A*′: convolution of *A* with *F*

## References

Ackman JB, Burbridge TJ, Crair MC (2012) Retinal waves coordinate patterned activity throughout the developing visual system. Nature 490:219–225.

Blankenship AG, Hamby AM, Firl A, Vyas S, Maxeiner S, Willecke K, Feller MB (2011) The role of neuronal connexins 36 and 45 in shaping spontaneous firing patterns in the developing retina. J Neurosci 31:9998–10008.

Butts DA, Kanold PO, Shatz CJ (2007) A burst-based Hebbian learning rule at retinogeniculate synapses links retinal waves to activity-dependent refinement. Plos Biol 5:651–661.

Chiappalone M, Bove M, Vato A, Tedesco M, Martinoia S (2006) Dissociated cortical networks show spontaneously correlated activity patterns during in vitro development. Brain Research 1093:41–53.

Cressie NAC (1993) Statistics for Spatial Data Wiley, New York.

Dayan P, Abbott LF (2001) Theoretical neuroscience: computational and mathematical modelling of neural systems MIT press, Cambridge, Massachusetts.

De la Rocha J, Doiron B, Shea-Brown E, Josíc K, Reyes A (2007) Correlation between neural spike trains increases with firing rate. Nature 448:802–806.

Dehorter N, Vinay L, Hammond C, Ben-Ari Y (2012) Timing of developmental sequences in different brain structures: physiological and pathological implications. Eur J Neurosci 35:1846–1856.

Demas J, Eglen SJ, Wong ROL (2003) Developmental loss of synchronous spontaneous activity in the mouse retina is independent of visual experience. J Neurosci 23:2851–2860.

Demas J, Sagdullaev BT, Green E, Jaubert-Miazza L, McCall MA, Gregg RG, Wong ROL, Guido Wet al. (2006) Failure to maintain eye-specific segregation in nob, a mutant with abnormally patterned retinal activity. Neuron 50:247–259.

Eggermont JJ (2010) Pair-correlation in the time and frequency domain. In Grun S, Rotter S, editors, Analysis of parallel spike trains, Vol. 7 of Springer series in Computational Neuroscience, pp. 77–102. Springer.

Eglen SJ, Weeks M, Jessop M, Simonotto J, Jackson T, Sernagor E (2014) A data repository and analysis framework for spontaneous neural activity recordings in developing retina. GigaScience 3:3.

Eytan D, Minerbi A, Ziv N, Marom S (2004) Dopamine induced dispersion of correlations between action potentials in networks of cortical neurons. J Neurophysiol 92:1817–1824.

Feller MB (2009) Retinal waves are likely to instruct the formation of eye-specific retinogeniculate projections. Neural Dev 4:24.

Godfrey KB, Eglen SJ (2009) Theoretical models of spontaneous activity generation and propagation in the developing retina. Mol Biosyst 5:1527–1535.

Grun S, Diesmann M, Aertsen A (2002) Unitary events in multiple single-neuron spiking activity: I. Detection and significance. Neural Computation 14:43–80.

Harris KD, Csicsvari J, Hirase H, Dragoi G, Buzsáki G (2003) Organisation of cell assemblies in the hippocampus. Nature 424:552–556.

Hunter JD, Milton JG (2003) Amplitude and frequency dependence of spike timing: implications for dynamic regulation. J Neurophysiol 90:387–394.

Isham V (1985) Marked point processes and their correlations. In Droesbeke F, editor, Spatial Processes and Spatial Time Series Analysis, pp. 63–75. Publications des Facultes Universitaires Saint-Louis, Brussels.

Jimbo Y, Tateno T, Robinson HPC (1999) Simultaneous induction of pathway-specific potentiation and depression in networks of cortical neurons. J Biophys 76:670–678.

Joris PX, Louage DH, Cardoen L, van der Heijden M (2006) Correlation index: a new metric to quantify temporal coding. Hearing research 216:19–30.

Kerschensteiner D, Wong ROL (2008) A precisely timed asynchronous pattern of ON and OFF retinal ganglion cell activity during propagation of retinal waves. Neuron 58:851–858.

Kirkby L, Sack GS, Firl A, Feller MB (2013) A role for correlated spontaneous activity in the assembly of nerual circuits. Neuron 80:1129–1144.

Kreuz T, Chicharro D, Houghton C, Andrzejak RG, Mormann F (2013) Monitoring spike train synchrony. J Neurophysiol 109:1457–1472.

Kreuz T, Haas JS, Morelli A, Abarbanel HDI, Politi A (2007a) Measuring spike train synchrony. J Neurosci Methods 165:151–161.

Kreuz T, Mormann F, Andrzejak RG, Kraskov A, Lehnertz K, Grassberger P (2007b) Measuring synchronisation in coupled model systems: A comparison of different approaches. Phys D 225:29–42.

Kruskal PB, Stanis JJ, McNaughton BL, Thomas PJ (2007) A binless correlation measure reduces the variability of memory reactivation estimates. Stat Med 26:3997–4008.

Lee CW, Eglen SJ, Wong ROL (2002) Segregation of ON and OFF retinogeniculate connectivity directed by patterned spontaneous activity. J Neurophysiol 88:2311–21.

Lee J, Lisberger SG (2013) Gamma synchrony predicts neuron-neuron correlations and correlations with motor behaviour in extrastriate visual area MT. J Neurosci 33:19677–88.

Li W (1990) Mutual information functions versus correlation functions. J. Stat. Phys. 60:823–837.

Maccione A, Hennig MH, Gandolfo M, Muthmann O, van Coppenhagen J, Eglen SJ, Berdondini L, Sernagor E (2014) Following the ontogeny of retinal waves: pan-retinal recordings of population dynamics in the neonatal mouse. J. Physiol 592:1545–1563.

Macke JH, Berens P, Ecker AS, Tolias AS, Bethge M (2009) Generating spike trains with specified correlation coefficients. Neural Comput 21:397–423.

MacLaren EJ, Charlesworth P, Coba MP, Grant SG (2011) Knockdown of mental disorder susceptibility genes disrupts neuronal network physiology in vitro. Mol Cell Neurosci 47:93–99.

McLaughlin T, Torborg CL, Feller MB, O’Leary DD (2003) Retinotopic map refinement requires spontaneous retinal waves during a brief critical period of development. Neuron 40:1147–1160.

Nakahara H, Amari S (2002) Information-geometric measure for neural spikes. Neural Comput 14:2269–2316.

Okun M, Yger P, Marguet SL, Gerard-Mercier F, Benucci A, Katzner S, Busse L, Carandini M, Harris KD (2012) Population rate dynamics and multineuron firing patterns in sensory cortex. J Neurosci 32:17108–19.

Parise CV, Harrar V, Ernst MO, Spence C (2013) Cross-correlation between auditory and visual signals promotes multisensory integration. Multisens Res 26:307–316.

Pasquale V, Massobrio P, Bologna LL, Chiappalone M, Martinoia S (2008) Self-organization and neuronal avalanches in networks of dissociated cortical neurons. Neuroscience 153:1354–1369.

Personius KE, Balice-Gordon RJ (2001) Loss of correlated motor neuron activity during synaptic competition at developing neuromuscular synapses. Neuron 31:395–408.

Personius KE, Chang Q, Mentis GZ, O’Donovan MJ, Balice-Gordon RJ (2007) Reduced gap junctional coupling leads to uncorrelated motor neuron firing and precocious neuromuscular synapse elimination. Proc Natl Acad Sci USA 104:11808–13.

Ripley BD (1976) The second-order analysis of stationary point processes. J Applied Probability 13:255–266.

Schlather M, Ribeiro PJ, Diggle PJ (2004) Detecting dependence between marks and locations of marked point processes. J R Stat Soc B 66:79–93.

Schlens J, Field GD, Gauthier JL, Grivich MI, Petrusca D, Sher A, Litke AM, Chichilnisky EJ (2006) The structure of multi-neuron firing patterns in primate retina. J Neurosci 26:8254–8266.

Schneidman E, Berry MJ, Segev R, Bialek W (2006) Weak pairwise correlations imply strongly correlated network states in a neural correlation. Nature 440:1007–1012.

Schreiber S, Fellous JM, Whitmer D, Tiesinga P, Sejnowski TJ (2003) A new correlation-based measure of spike timing reliability. Neurocomputing 52:925–931.

Shi X, Lu Q (2009) Burst synchronization of electrically and chemically coupled map-based neurons. Phys A 388:2410–2419.

Simpson EH (1949) Measurement of diversity. Nature 163:688

Stafford BK, Sher A, Litke AM, Feldheim DA (2009) Spatial-temporal patterns of retinal waves underlying activity-dependent refinement of retinofugal projections. Neuron 64:200–212.

Stoyan D (1984) Correlations of the marks of marked point processes - statistical inference and simple models. Elektron. Informationsverarbeitung und Kybernetik 20:285–294.

Stoyan D, Stoyan H (1994) Fractals, random shapes and point fields John Wiley and Sons, Chichester.

Sun C, Warland DK, Ballesteros JM, van der List D, Chalupa LM (2008) Retinal waves in mice lacking the beta2 subunit of the nicotinic acetylcholine receptor. Proc Natl Acad Sci USA 105:13638–43.

Van Rossum MC (2001) A novel spike distance. Neural Comput 13:751–763.

Victor JD, Purpura K (1997) Metric-space analysis of spike trains: theory, algorithms and application. Network 8:127–164.

Walters E, Segonds-Pinchon A, Nicol AU (2008) The correlation determinant in tests for synchronization in neuronal spike data. J Neurosci Methods 172:60–66.

Wong ROL (1999) Retinal waves and visual system development. Annu Rev Neurosci 22:29–47.

Wong ROL, Meister M, Shatz CJ (1993) Transient period of correlated bursting activity during development of the mammalian retina. Neuron 11:923–938.

Xu HP, Furman M, Mineur YS, Chen H, King SL, Zenisek D, Zhou ZJ, Butts DA, Tian N, Picciotto MR, Crair MC (2011) An instructive role for patterened spontaneous retinal activity in mouse visual map development. Neuron 70:1115–1127.

Zhang LI, Tao HW, Holt CE, Harris WA, Poo M (1998) A critical window for cooperation and competition among developing retinotectal synapses. Nature 395:37–44.

